# The FDA-approved drug nitazoxanide is a potent inhibitor of human seasonal coronaviruses acting at postentry level: effect on the viral spike glycoprotein

**DOI:** 10.1101/2022.07.13.499346

**Authors:** Sara Piacentini, Anna Riccio, Silvia Santopolo, Silvia Pauciullo, Simone La Frazia, Antonio Rossi, Jean-Francois Rossignol, M. Gabriella Santoro

**Author notes:** Correspondence and requests for materials should be addressed to M.G.S. These authors contributed equally to this work (Sara Piacentini, Anna Riccio).

## Abstract

*Coronaviridae* is recognized as one of the most rapidly evolving virus family as a consequence of the high genomic nucleotide substitution rates and recombination. The family comprises a large number of enveloped, positive-sense single-stranded RNA viruses, causing an array of diseases of varying severity in animals and humans. To date, seven human coronaviruses (HCoV) have been identified, namely HCoV-229E, HCoV-NL63, HCoV-OC43 and HCoV-HKU1, which are globally circulating in the human population (seasonal HCoV, sHCoV), and the highly pathogenic SARS-CoV, MERS-CoV and SARS-CoV-2. Seasonal HCoV are estimated to contribute to 15-30% of common cold cases in humans; although diseases are generally self-limiting, sHCoV can sometimes cause severe lower respiratory infections, as well as enteric and neurological diseases. No specific treatment is presently available for sHCoV infections. Herein we show that the anti-infective drug nitazoxanide has a potent antiviral activity against three human endemic coronaviruses, the Alpha-coronaviruses HCoV-229E and HCoV-NL63, and the Beta-coronavirus HCoV-OC43 in cell culture with IC_50_ ranging between 0.05 and 0.15 μg/ml and high selectivity indexes. We found that nitazoxanide does not affect HCoV adsorption, entry or uncoating, but acts at postentry level and interferes with the spike glycoprotein maturation, hampering its terminal glycosylation at an endoglycosidase H-sensitive stage. Altogether the results indicate that nitazoxanide, due to its broad-spectrum anti-coronavirus activity, may represent a readily available useful tool in the treatment of seasonal coronavirus infections.

## 1. Introduction

Coronaviruses (CoV), members of the family *Coronaviridae* (order *Nidovirales)*, comprise a large number of enveloped, positive-sense single-stranded RNA viruses causing respiratory, enteric, hepatic and neurological diseases of varying severity in animals and humans (Cui et al., 2019; Fung and Liu, 2019). Coronaviruses have the largest identified RNA genomes (typically ranging from 27 to 32 kb) containing multiple open reading frames with an invariant gene order [a large replicase-transcriptase gene preceding structural (S-E-M-N) and accessory genes] (Fung and Liu, 2019). On the basis of their phylogenetic relationships and genomic structures, CoV are subdivided in four genera: Alpha-, Beta-, Gamma- and Delta-coronavirus; among these Alpha- and Beta-CoVs infect only mammals, whereas Gamma- and Delta-CoVs infect birds, and only occasionally can infect mammals (Cui et al., 2019). Human coronaviruses (HCoV) were discovered in the 1960s and were originally thought to cause only mild disease in humans (Cui et al., 2019; Fung and Liu, 2019). This view changed in 2002 with the SARS (Severe Acute Respiratory Syndrome) epidemic and in 2012 with the MERS (Middle East Respiratory Syndrome) outbreak, two zoonotic infections that resulted in mortality rates greater than 10% and 35%, respectively (Cui et al., 2019; Fung and Liu, 2019). Near the end of 2019, the seventh coronavirus known to infect humans, SARS-CoV-2, phylogenetically in the SARS-CoV clade, emerged in Wuhan, China. SARS-CoV-2 turned out to be a far more serious threat to public health than SARS-CoV and MERS-CoV because of its ability to spread more efficiently, making it difficult to contain worldwide with more than 760 million confirmed cases and over 6.8 million deaths reported worldwide, as of March 20th, 2023 (https://covid19.who.int/). The clinical features of COVID-19, the disease associated with SARS-CoV-2, vary ranging from asymptomatic state to respiratory symptoms that, in a subset of patients, may progress to pneumonia, acute respiratory distress syndrome (ARDS), multi organ dysfunction and death (Lamers and Haagmans, 2022; Tang et al., 2020).

Only two HCoV, HCoV-OC43 and HCoV-229E, were known prior to the emergence of SARS-CoV (Hamre and Procknow, 1966; Tyrrel and Bynoe, 1965), while two more, HCoV-NL63 and HCoV-HKU1, were identified between 2004 and 2005 (van der Hoek et al., 2004; Woo et al., 2005). HCoV-OC43 and HCoV-HKU1 likely originated in rodents, while HCoV-229E and HCoV-NL63, similarly to SARS-CoV and MERS-CoV, originated in bats (Fung and Liu, 2019).

These four HCoV (seasonal HCoV, sHCoV) are globally distributed and are estimated to contribute to 15–30% of cases of common cold in humans (Lim et al., 2016; Liu et al., 2021). Although diseases are generally self-limiting, sHCoV can sometimes cause severe lower respiratory infections, including life-threatening pneumonia and bronchiolitis especially in infants, elderly people, or immunocompromised patients (Chiu et al., 2005; Gorse et al., 2009; Pene et al., 2003; Zhang et al., 2022); in addition, besides respiratory illnesses, sHCoV may cause enteric and neurological diseases (Arbour et al., 2000; Jacomy et al., 2006; Morfopoulou et al., 2016; Risku et al., 2010), while a possible involvement of HCoV-229E in the development of Kawasaki disease was suggested (Esper et al., 2005; Shirato et al., 2014).

Whereas all seasonal coronaviruses cause respiratory tract infections, HCoV-OC43, HCoV-229E, HCoV-NL63 and HCoV-HKU1 are genetically dissimilar (Fig. 1A), belonging to two distinct taxonomic genera (Alpha and Beta), and use different receptors that represent the major determinants of tissue tropism and host range (Fung and Liu, 2019). HCoV-229E and HCoV-NL63 have adopted cell surface enzymes as receptors, such as aminopeptidase N (APN) for HCoV-229E and angiotensin converting enzyme 2 (ACE2) for HCoV-NL63, while HCoV-OC43 and HCoV-HKU1 use 9-*O*-acetylated sialic acid as a receptor (Fung and Liu, 2019; Lim et al., 2016). In all cases, sHCoV infection is initiated by the binding of the spike (S) glycoprotein, anchored into the viral envelope, to the host receptor (Fung and Liu, 2019).

**Figure 1.**
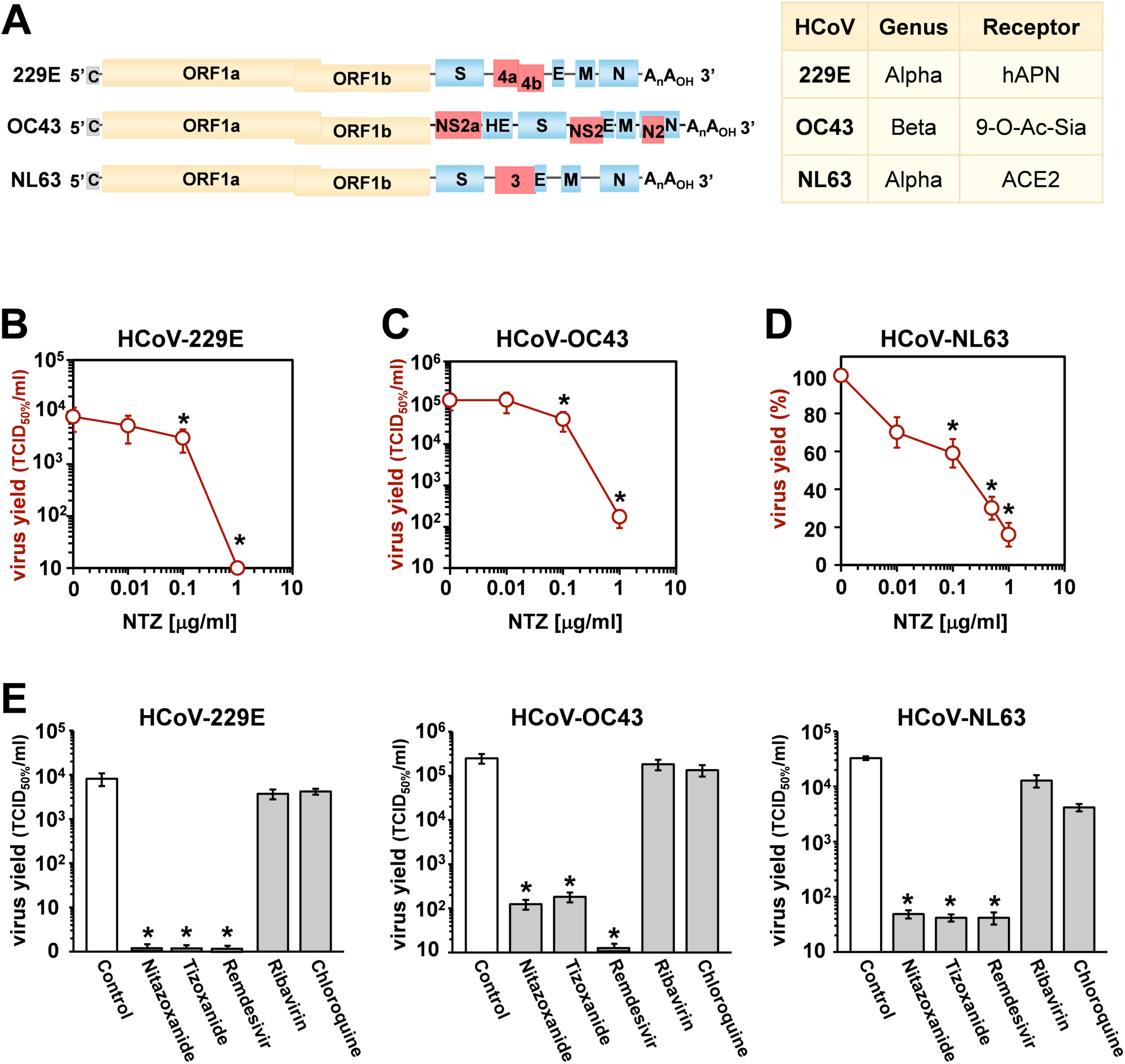
Antiviral activity of nitazoxanide against human seasonal coronaviruses. (**A**) Schematic representation of genome structure, classification and receptors of the human coronaviruses HCoV-229E, HCoV-OC43 and HCoV-NL63. ORF1a and ORF1b are represented as yellow boxes; genes encoding structural proteins spike (S), nucleocapsid (N), envelope (E), membrane (M), and hemagglutinin-esterase (HE) are shown as blue boxes, and genes encoding accessory proteins are shown as red boxes. hAPN, human aminopeptidase N; 9-O-Ac-Sia, N-acetyl-9-O-acetylneuraminic acid; ACE2, angiotensin-converting enzyme 2. (**B-D**) MRC-5 (B,C) and LLC-MK2 (D) cells mock-infected or infected with HCoV-229E (B), HCoV-OC43 (C) or HCoV-NL63 (D) at an MOI of 0.1 TCID_50_/cell were treated with different concentrations of NTZ or vehicle immediately after the adsorption period. In the case of HCoV-NL63, NTZ was removed at 48h after infection. Virus yield in cell supernatants was determined at 48 (B), 96 (C) or 120 (D) hours p.i. by infectivity assay (B,C) or RNA quantification by qRT-PCR (D). Data, expressed as TCID_50_/ml (B,C) or percent of untreated control (D), represent the mean ± S.D. of duplicate samples. (**E**) MRC-5 cells infected with HCoV-229E or HCoV-OC43 and LLC-MK2 cells infected with HCoV-NL63 (0.1 TCID_50_/cell) were treated with 3 µM nitazoxanide, the NTZ active metabolite tizoxanide, remdesivir, chloroquine and ribavirin after virus adsorption. In the case of HCoV-NL63, NTZ and tizoxanide were removed at 48h after infection. Virus yields were determined at 48 (HCoV-229E), 72 (HCoV-OC43) or 120 (HCoV-NL63) hours p.i. by infectivity assay. Data, expressed as TCID_50_/ml, represent the mean ± S.D. of duplicate samples. *= p< 0.05; ANOVA test.

The coronavirus spike protein is a trimeric class-I fusion glycoprotein (Li, 2016); each monomer, with a molecular weight of 150–200 kDa after N-linked glycosylation (Lim et al., 2016), is synthesized as a fusogenically-inactive precursor that assembles into an inactive homotrimer, which is endoproteolytically cleaved by cellular proteases giving rise to a metastable complex of two functional subunits: S1 (bulb) containing the receptor-binding domain responsible for recognition and attachment to the host receptor, and the membrane-anchored S2 (stalk) that contains the fusion machinery (Li, 2016; Santopolo et al., 2021a). During synthesis in the infected cell, the nascent spike is imported into the endoplasmic reticulum (ER), where the protein is glycosylated (Santopolo et al., 2021a; Sicari et al., 2020). S glycoproteins passing the quality control mechanisms of the ER are transported to the ER/Golgi intermediate compartment (ERGIC), the presumed site of viral budding (Fung and Liu, 2019; Li, 2016).

Despite the fact that endemic seasonal coronaviruses may cause severe, life-threatening diseases in a subset of patients, no specific treatment is available for HCoV infections.

Nitazoxanide, a thiazolide originally developed as an antiprotozoal agent and used in clinical practice for the treatment of infectious gastroenteritis (Rossignol et al., 2001, 2006), and second-generation thiazolides have emerged as a new class of broad-spectrum antiviral drugs (Rossignol, 2014). Herein we investigated the antiviral activity of nitazoxanide against three human endemic coronaviruses belonging to two different genera: the Alpha-coronaviruses HCoV-229E and HCoV-NL63, and the Beta-coronavirus HCoV-OC43. We report that nitazoxanide is a potent inhibitor of HCoV replication acting at postentry level and interfering with Alpha- and Beta- sHCoV spike glycoprotein maturation.

## 2. Materials and methods

### 2.1 Cell culture and treatments

Human normal lung MRC-5 fibroblasts (American Type Culture Collection, ATCC, CCL-171) and rhesus monkey kidney LLC-MK2 cells (a kind gift from Lia van der Hoek, Academic Medical Center, University of Amsterdam) were grown at 37°C in a 5% CO_2_ atmosphere in minimal essential medium (MEM, Gibco 32360-026) for LLC-MK2 cells or EMEM (ATCC 30-2003) for MRC-5 cells, supplemented with 10% fetal calf serum (FCS), 2 mM glutamine and antibiotics. Nitazoxanide [2-acetyloxy-*N*-(5-nitro-2-thiazolyl) benzamide, Alinia] (NTZ) and tizoxanide (TIZ) (Romark Laboratories, L.C.), remdesivir (MedChemExpress), ribavirin and chloroquine (Sigma-Aldrich), dissolved respectively in DMSO stock solution (NTZ, TIZ, remdesivir) or water (ribavirin, chloroquine), were diluted in culture medium, added to infected cells after the virus adsorption period, and maintained in the medium for the duration of the experiment, unless differently specified. Controls received equal amounts of DMSO vehicle, which did not affect cell viability or virus replication. Cell viability was determined by the 3-(4,5-dimethylthiazol-2-yl)-2,5-diphenyltetrazolium bromide (MTT) to MTT formazan conversion assay (Sigma-Aldrich), as described (Pizzato Scomazzon et al., 2019). The 50% lethal dose (LD_50_) was calculated using Prism 5.0 software (Graph-Pad Software Inc.). Microscopical examination of mock-infected or virus-infected cells was performed daily to detect virus-induced cytopathic effect and possible morphological changes and/or cytoprotection induced by the drug. Microscopy studies were performed using a Leica DM-IL microscope and images were captured on a Leica DC 300 camera using Leica Image-Manager500 software.

### 2.2 Coronavirus infection and titration

Human coronaviruses HCoV-229E (ATCC), HCoV-OC43 (ATCC) and HCoV-NL63 (strain Amsterdam-1, a kind gift from Lia van der Hoek, University of Amsterdam), were used for this study. Due to its poor ability to grow in cell culture the fourth known seasonal HCoV, HKU1, was not investigated. For virus infection, confluent MRC-5 (229E and OC43) or LLC-MK2 (NL63) cell monolayers were infected with HCoV for 1 (229E and OC43) or 2 (NL63) hours at 33°C at a multiplicity of infection (MOI) of 0.1 or 0.5 TCID_50_ (50% tissue culture infectious dose)/cell. After the adsorption period, the viral inoculum was removed, and cell monolayers were washed three times with phosphate-buffered saline (PBS). Cells were maintained at 33°C in growth medium containing 2% FCS. Virus yield was determined at different times after infection by TCID_50_ infectivity assay, as described previously (Santoro et al., 1988). The IC_50_ (50% inhibitory concentration) of the compounds tested was calculated using Prism 5.0 software.

### 2.3 HCoV RNA extraction and quantification

Measurement of sHCoV genomic RNA was performed by real-time quantitative reverse transcription-PCR (qRT-PCR), as described (Coccia et al., 2017). Briefly, total RNA from mock-infected or HCoV-infected cells was prepared using ReliaPrep RNA Cell Miniprep System (Promega) and reverse transcription was performed with PrimeScript RT Reagent Kit (Takara) according to the manufacturer’s protocol. Extracellular viral RNA was extracted from 200 µl of the supernatant of mock-infected or sHCoV-infected cell cultures with Viral Nucleic Acid Extraction Kit II (Geneaid) and 10 µl were subjected to reverse transcription using SuperScript™ VILO™ cDNA Synthesis Kit (Life Technologies) as described in the manufacturer’s protocol. Real-time PCR analysis was performed with CFX-96 (Bio-Rad) using SensiFAST SYBR® kit (Bioline) and primers specific for the membrane protein gene of HCoV-OC43 and HCoV-229E (Vijgen et al., 2005) or for the nucleoprotein gene of HCoV-NL63 (Pyrc et al., 2006). Relative quantities of selected mRNAs were normalized to ribosomal L34 RNA levels in the same sample. The sequences of the L34 primers were as follows: sense 5′-GGCCCTGCTGACATGTTTCTT-3′, antisense 5′-GTCCCGAACCCCTGGTAATAGA-3′. All reactions were made in triplicate using samples derived from three biological repeats.

### 2.4 HCoV genomic RNA transfection

For sHCoV genomic RNA transfection experiments, MRC-5 cell monolayers were infected with HCoV-OC43 or HCoV-229E for 1h at 33°C at an MOI of 0.1 TCID_50_/cell and sHCoV genomic RNA was extracted from the supernatants at 24h p.i. using TRIzol-LS reagent (Life Technologies) as described in the manufacturer protocol. MRC-5 cell monolayers were mock-transfected or transfected with sHCoV genomic RNA (1 μg/ml) using TransIT-mRNA Transfection Kit (Mirus Bio) at 33°C. After 4h, transfection medium was removed and cells were treated with NTZ or vehicle and maintained at 33°C in growth medium containing 2% FCS. At 24h after treatment, culture supernatants were collected for virus progeny titer determination, and cell monolayers were processed for viral proteins detection.

### 2.5 Protein Analysis, Western blot and endoglycosidase digestion

For analysis of proteins whole-cell extracts (WCE) were prepared after lysis in High Salt Buffer (HSB) (Santopolo et al., 2021b). Briefly, cells were washed twice with ice-cold PBS and then lysed in HSB (40 µl). After one cycle of freeze and thaw, and centrifugation at 16,000×g (10 min at 4°C), supernatant and pellet fractions were collected (Santoro et al., 1982). For Western blot analysis, cell extracts (20 µg/sample) were separated by SDS-PAGE under reducing conditions and blotted to a nitrocellulose membrane. Membranes were incubated with rabbit polyclonal anti-HCoV-OC43 spike (CSB-PA336163EA01HIY, Cusabio) or nucleocapsid (40643-T62, Sino Biological) antibodies, anti-HCoV-229E spike (PAB21477-100, LGC NAC Company) or nucleocapsid (40640-T62, Sino Biological) antibodies, anti β-actin (A2066, Sigma-Aldrich) antibodies, and monoclonal α-tubulin (T5168, Sigma-Aldrich) antibodies, followed by decoration with peroxidase-labeled anti-rabbit or anti-mouse IgG (Super-Signal detection kit, Pierce). Spike proteins densitometric and molecular weight analysis were performed with Image Lab Software 6.1, after acquisition on a ChemiDoc XRS+ (Bio-Rad).

For endoglycosidase digestion experiments, samples containing the same amount of protein (20 µg/sample) were processed for endoglycosidase-H (Endo-H, NEB) digestion using 5 milliunits Endo-H for 16h at 37°C, according to the manufacturer’s protocol (Riccio et al., 2022). Samples were separated by SDS-PAGE and blotted as described above. Quantitative evaluation of proteins was determined as described (Amici et al., 2015; Santoro et al., 1989). All results shown are representative of at least three independent experiments.

### 2.6 Immunofluorescence microscopy

MRC-5 cells infected with HCoV-OC43 or HCoV-229E were grown in 8-well chamber slides (Lab-Tek II) and, after the adsorption period, were treated with NTZ, or vehicle for 24h. Cells were fixed, permeabilized and processed for immunofluorescence as described (Riccio et al., 2022) using anti-HCoV-OC43 or anti-HCoV-229E spike antibodies, followed by decoration with Alexa Fluor 555-conjugated antibodies (Molecular Probes, Invitrogen). Nuclei were stained with Hoechst 33342 (Molecular Probes, Invitrogen). Images were captured using a ZEISS Axio Observer Inverted Microscope and analyzed using ZEN 3.1 (blue edition) software. For confocal microscopy, images were acquired on Olympus FluoView FV-1000 confocal laser scanning system (Olympus America Inc., Center Valley, PA) and analyzed using Imaris (v6.2) software (Bitplane, Zurich, Switzerland). Images shown in all figures are representative of at least three random fields (scale-bars are indicated).

### 2.7 Statistical analysis

Statistical analyses were performed using Prism 5.0 software (GraphPad Software). Comparisons between two groups were made using Student’s *t*-test; comparisons among groups were performed by one-way ANOVA with Bonferroni adjustments. *p* values ≤0.05 were considered significant. Data are expressed as the means ± standard deviations (SD) of results from duplicate or quadruplicate samples. Each experiment (in duplicate) was repeated at least twice.

## 3. Results

### 3.1 Antiviral activity of nitazoxanide against human seasonal coronaviruses

Three different human globally distributed coronaviruses, HCoV-229E, HCoV-OC43, and HCoV-NL63, were utilized for the current study. The genomic structure, classification, and receptors of these HCoVs are summarized in Fig. 1A.

Nitazoxanide antiviral activity was first investigated in human lung MRC-5 and monkey kidney LLC-MK2 cells infected with HCoVs 229E (Fig. 1B), OC43 (Fig. 1C) and NL63 (Fig. 1D) at the MOI of 0.1 TCID_50_/cell, and treated with different concentrations of the drug starting after the virus adsorption period. In the case of HCoV-NL63, which required 120 hours for infectious progeny production, nitazoxanide was removed at 48h after infection. At 48 (229E), 96 (OC43) and 120 (NL63) hours after infection, viral titers were determined in the supernatant of infected cells by TCID_50_ assay (Fig. 1B,C) or viral RNA quantification (Fig. 1D; Supplementary Fig. 1A,B). Nitazoxanide showed a remarkable antiviral activity against all three HCoVs, reducing virus yield dose-dependently with IC_50_ values in the submicromolar range.

To compare the effect of nitazoxanide with other antiviral drugs, MRC-5 and LLC-MK2 cells were infected with the different HCoVs at the MOI of 0.1 TCID_50_/cell and treated with nitazoxanide, the NTZ bioactive metabolite tizoxanide, the antiviral drugs remdesivir and ribavirin, or chloroquine at the same concentration (3 μM) starting after virus adsorption. In the case of HCoV-NL63, NTZ and tizoxanide were removed at 48h after infection. Virus yields were determined at 48 (HCoV-229E), 72 (HCoV-OC43) or 120 (HCoV-NL63) hours after infection by TCID_50_ assay. Nitazoxanide, tizoxanide and remdesivir showed comparable antiviral activity against all three HCoV, whereas chloroquine and ribavirin were not found to be effective when treatment was started after infection (Fig. 1E).

Next, the effect of short-treatment with nitazoxanide was investigated in MRC-5 cells infected with HCoV-OC43 and HCoV-229E. MRC-5 cells were infected with the OC43 or 229E HCoV strains at the MOI of 0.5 TCID_50_/cell and treated with different concentrations of the drug starting after the virus adsorption period. At 24 hours after infection, viral titers were determined in the supernatant of infected cells by TCID_50_ assay (Fig. 2A) and viral RNA level quantification (Supplementary Fig. 1C,D); in parallel, the effect of nitazoxanide on mock-infected cell viability was determined by MTT assay. The results, shown in Fig. 2A,B, confirmed a potent antiviral activity of nitazoxanide, with IC_50_ values of 0.15 μg/ml and 0.05 μg/ml and selectivity indexes higher than 330 and 1000 for HCoV-OC43 and HCoV-229E respectively.

**Figure 2.**
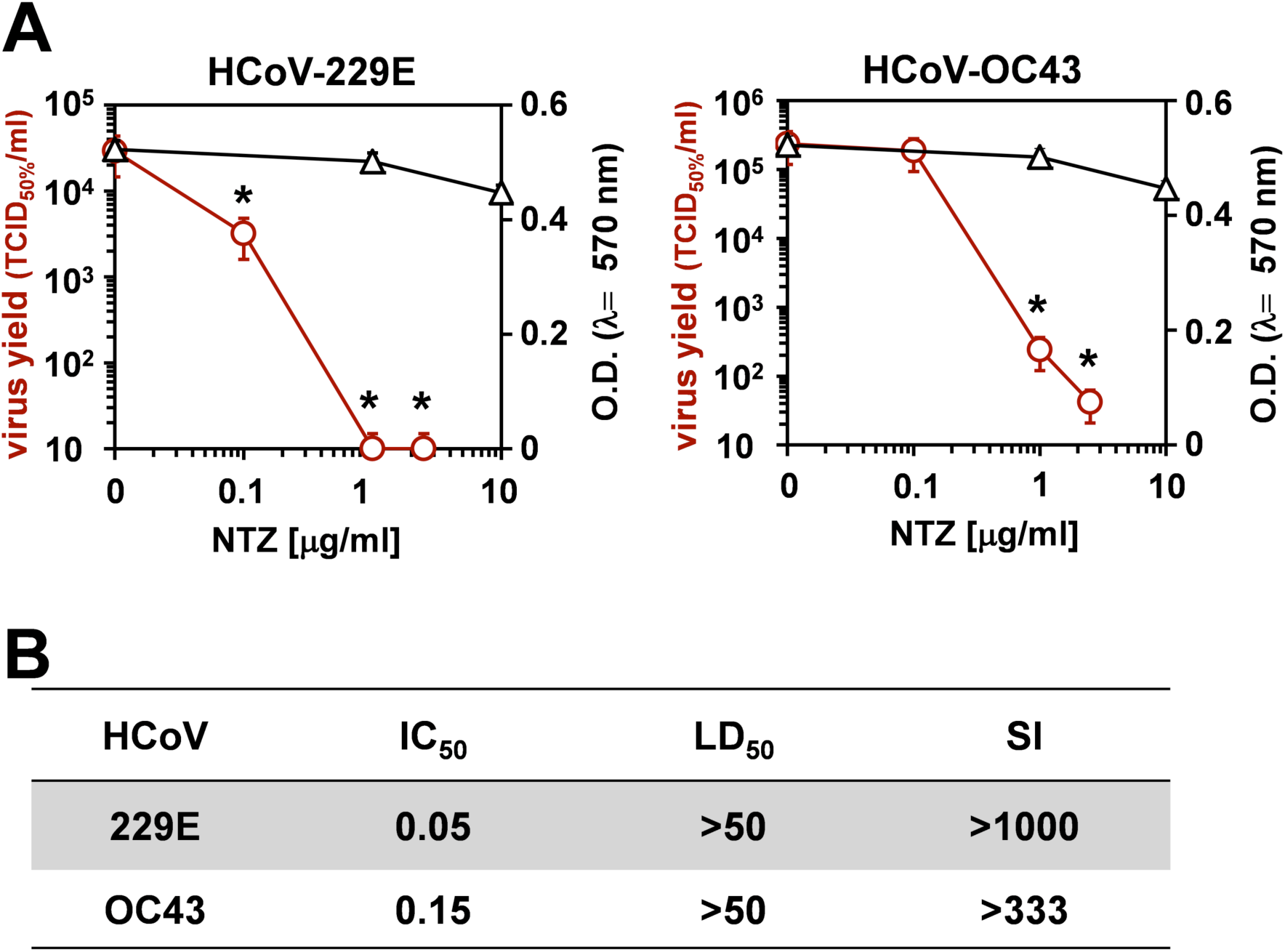
Effect of short treatment with nitazoxanide in OC43 and 229E HCoV-infected human lung cells. (**A,B**) MRC-5 cell monolayers mock-infected or infected with HCoV-229E and HCoV-OC43 (0.5 TCID_50_/cell) were treated with different concentrations of NTZ or vehicle immediately after the adsorption period. Virus yield (Ο) was determined at 24h p.i. by infectivity assay. In parallel, cell viability (△) was determined by MTT assay in mock-infected cells (A). Absorbance (O.D.) of converted dye was measured at λ = 570nm. IC_50_ and LD_50_ in μg/ml (B) were calculated using Prism 5.0 software. Data represent the mean ± S.D. of duplicate samples. Selectivity Indexes (SI) are indicated. *= p< 0.05; ANOVA test.

### 3.2 Nitazoxanide acts at postentry level

To determine the effect of NTZ treatment before virus infection, MRC-5 cells were treated with NTZ (1 or 2.5 μg/ml) or vehicle for 2h and the drug was removed before infection with OC43 or 229E HCoVs (0.5 TCID_50_/cell). In parallel, MRC-5 cells were infected with OC43 or 229E HCoVs (0.5 TCID_50_/cell) in the absence of the drug, and treated with 1 or 2.5 μg/ml NTZ or vehicle starting immediately after the adsorption period for the duration of the experiment. Alternatively, MRC-5 cells were treated 2h before infection and treatment was continued during and after the adsorption period. Virus yield was determined at 24h p.i. by TCID_50_ assay. As shown in Fig. 3A,B, NTZ pretreatment did not significantly affect HCoV replication when the drug was removed before infection; on the other hand, treatment started after infection was effective in reducing infectious progeny production by approximately 3-fold in both HCoV models, without affecting cell viability (LD_50_> 50 μg/ml).

**Figure 3.**
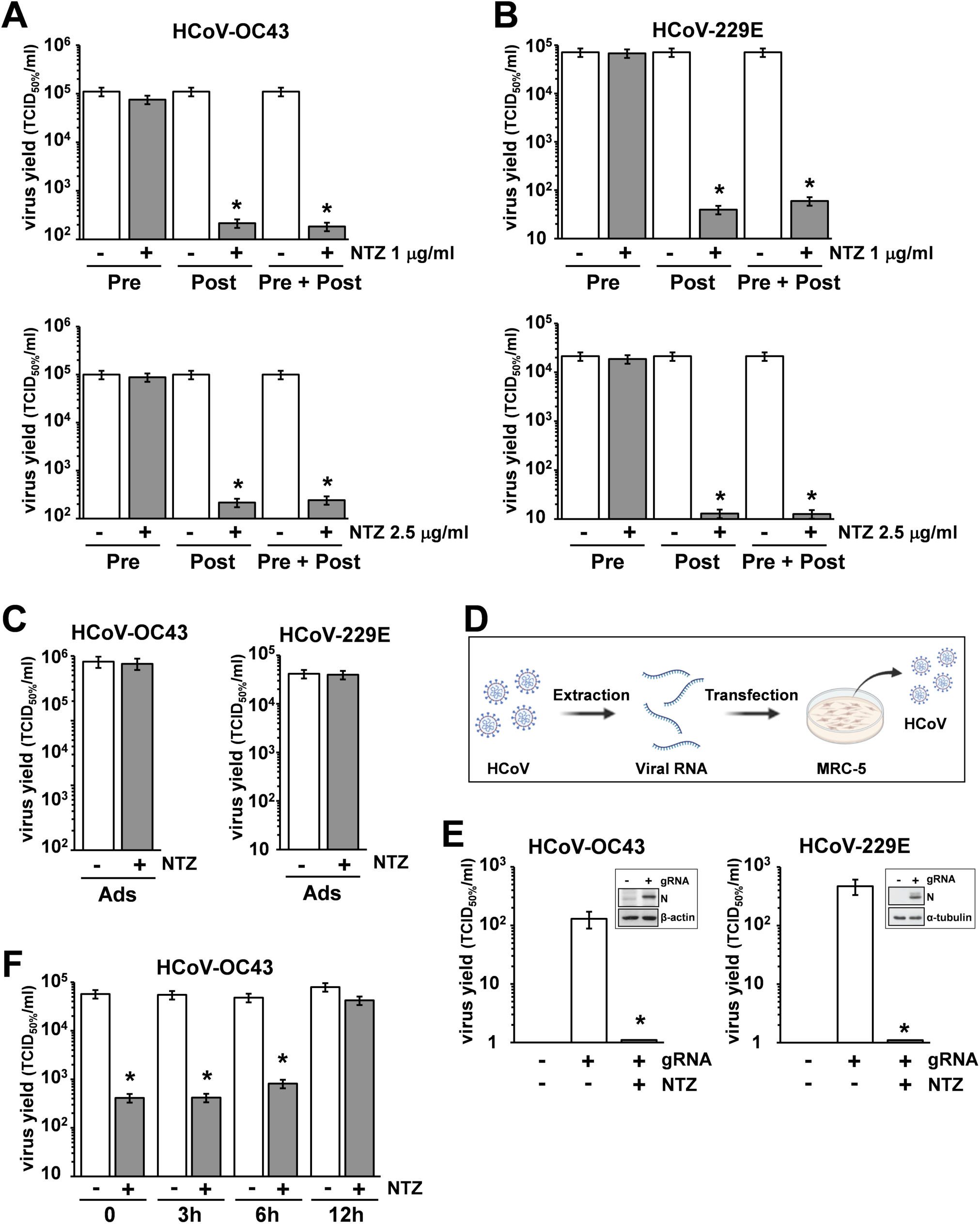
Nitazoxanide acts at postentry level. (**A,B**) MRC-5 cells mock-infected or infected with HCoV-OC43 (A) or HCoV-229E (B) (0.5 TCID_50_/cell) were treated with NTZ 1 µg/ml (A,B, top; filled bars), NTZ 2.5 µg/ml (A,B, bottom; filled bars) or vehicle (empty bars) 2h before infection (Pre), after the adsorption period (Post), or 2h before infection and treatment was continued during and after the adsorption period (Pre + Post). Virus yield was determined at 24h p.i. by TCID_50_ infectivity assay. (**C**) MRC-5 cells mock-infected or infected with HCoV-OC43 or HCoV-229E were treated with NTZ (1 µg/ml) or vehicle only during the adsorption period (Ads). Virus yield was determined at 24h p.i. by infectivity assay. (**A-C**) Data represent the mean ± S.D. of duplicate samples. *= p< 0.01; Student’s *t*-test. (**D**) Schematic representation of HCoV genomic RNA transfection assay. (**E**) MRC-5 cells were transfected with HCoV-OC43 or HCoV-229E genomic RNA for 4h and treated with NTZ (2.5 µg/ml) or vehicle for 24h. Virus yield was determined at 24h after treatment in the supernatant of transfected cells by TCID_50_ infectivity assay. Nucleocapsid (N) protein levels in HCoV-OC43 or HCoV-229E RNA-transfected cells are shown (insets). (**F**) MRC-5 cells infected with HCoV-OC43 were treated with NTZ (1 µg/ml) or vehicle at the indicated times after infection. Virus yield was determined at 24h p.i. by infectivity assay. (**E-F**) Data represent the mean ± S.D. of duplicate samples. *= p< 0.01; Student’s *t*-test.

In a different experiment, MRC-5 cells were infected with OC43 or 229E HCoV (0.5 TCID_50_/cell) and treated with NTZ (1 μg/ml) or vehicle only during the 1h adsorption period, after which time the drug was removed. Virus yield was determined at 24h p.i. by TCID_50_ assay. As shown in Fig. 3C, NTZ treatment during virus adsorption did not affect HCoV replication. These results suggest that nitazoxanide acts at postentry level.

In order to rule out any effect of the drug on virus adsorption, entry or uncoating, OC43 and 229E HCoV genomic RNA was extracted and transfected into MRC-5 monolayers, as described in Materials and Methods, and schematically represented in Fig. 3D. After 4h, the transfection medium was removed and cells were treated with NTZ (2.5 µg/ml) or vehicle for 24h. Virus yield was determined at 24h after treatment in the supernatant of transfected cells. As shown in Fig. 3E, genomic RNA transfection resulted in the production of infectious viral progeny (10^2^-10^3^TCID_50_/ml) already at 24h after transfection. Higher virus titers were detected at later times after transfection (data not shown). NTZ treatment greatly reduced the production of both HCoV-OC43 and HCoV-229E infectious particles after RNA transfection. Altogether, these results demonstrate that nitazoxanide is not acting on HCoV adsorption, entry or uncoating.

### 3.3 Nitazoxanide treatment impairs HCoV-OC43 and HCoV-229E spike maturation

Interestingly, NTZ-treatment initiated between 0 and 6h p.i. was equally effective in inhibiting HCoV-OC43 virus replication, whereas the antiviral activity was impaired when treatment was started as late as 12h p.i. (Fig. 3F). Similar results were obtained after HCoV-229E infection (Supplementary Fig. 2).

To investigate whether NTZ may affect human coronavirus structural proteins expression, MRC-5 cells were infected with HCoV-OC43 (0.5 TCID_50_/cell) and treated with 0.1, 1 or 2.5 μg/ml NTZ or vehicle starting immediately after the adsorption period. At 24h p.i., levels of the viral nucleocapsid (N) and spike (S) proteins were determined in the infected cells by immunoblot analysis using specific antibodies, and virus yield was determined in the culture supernatants by infectivity assay. As shown in Fig. 4A, no significant differences in N and S protein levels were detected in NTZ-treated cells, as compared to control, at concentrations that caused a >99% reduction in viral yield in the same samples (Fig. 4B). Comparable levels of S-protein were also detected by confocal microscopy in MRC-5 cells infected with OC43 or 229E HCoVs (0.5 TCID_50_/cell) and treated with NTZ (1 μg/ml) or vehicle after the adsorption period for 24h (Fig. 4C; Supplementary Fig. 3). Interestingly, treatment with NTZ at concentrations higher than 0.1 μg/ml caused an evident alteration in the electrophoretic mobility pattern of the spike glycoprotein (Fig. 4A). An alteration in the molecular mass of the HCoV-OC43 spike protein of approximately 5.8 kDa was in fact detected in MRC-5 cells treated with 2.5 μg/ml NTZ (Fig. 5A,C). In a parallel experiment, a similar alteration in the HCoV-229E spike molecular mass (8.3 kDa) was detected in MRC-5 cells treated with 2.5 μg/ml NTZ (Fig. 5B,D), indicating an effect of the drug in the glycoprotein maturation process. These results are in line with our previous observation that the thiazolide affects the maturation of the SARS-CoV-2 S glycoprotein in human cells transfected with plasmids encoding different SARS-CoV-2 spike variants (Riccio et al., 2022). In this case, we have found that nitazoxanide blocks the maturation of the SARS-CoV-2 S glycoprotein at a stage preceding resistance to digestion by endoglycosidase-H (Endo-H), an enzyme that removes *N*-linked carbohydrate chains that have not been terminally glycosylated (Fig. 5E) (Ohuchi et al., 1997), thus impairing SARS-CoV-2 S intracellular trafficking and infectivity.

**Figure 4.**
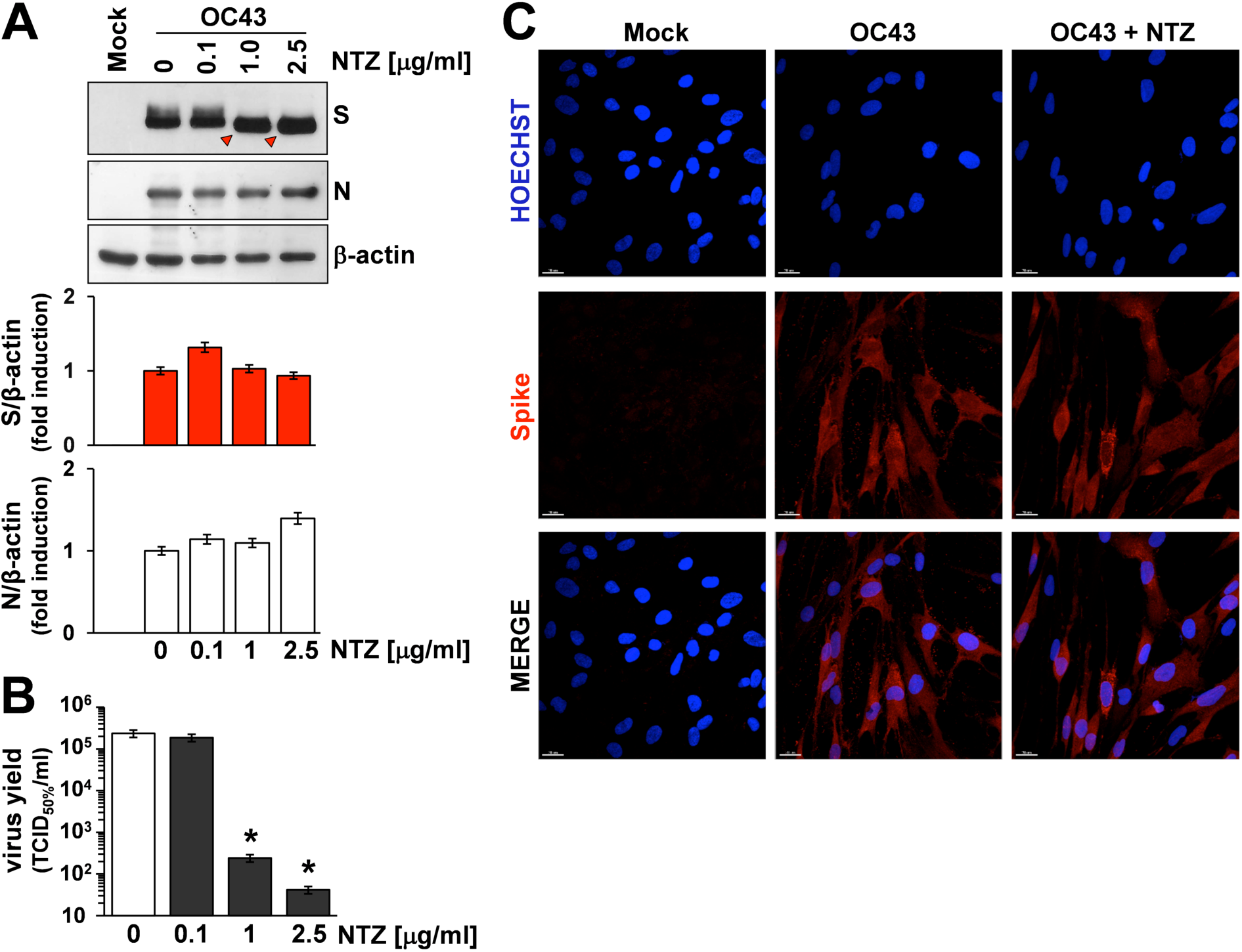
Effect of Nitazoxanide on HCoV-OC43 spike expression. (**A,B**) MRC-5 cells were mock-infected or infected with HCoV-OC43 at an MOI of 0.5 TCID_50_/cell and treated with different concentrations of NTZ or vehicle immediately after the virus adsorption period. After 24h, whole cell extracts (WCE) were analyzed for levels of viral S and N proteins by IB using anti-spike and anti-N HCoV-OC43 antibodies (A, top), and quantitated by scanning densitometry. Relative amounts of S and N proteins were determined after normalizing to β-actin (A, bottom). The faster-migrating form of the S protein in NTZ-treated cells is indicated by red arrowheads. In the same experiment virus yield was determined at 24h p.i. in the supernatant of infected cells by TCID_50_ infectivity assay (B). Data represent the mean ± S.D. of duplicate samples. *= p< 0.01; ANOVA test. **(C)** Confocal images of HCoV-OC43 spike glycoprotein (red) in MRC-5 cells mock-infected or infected with HCoV-OC43 at an MOI of 0.5 TCID_50_/cell and treated with NTZ (1 µg/ml) or vehicle for 24h. Nuclei are stained with Hoechst (blue). Merge images are shown. Scale bar, 20 µm.

**Figure 5.**
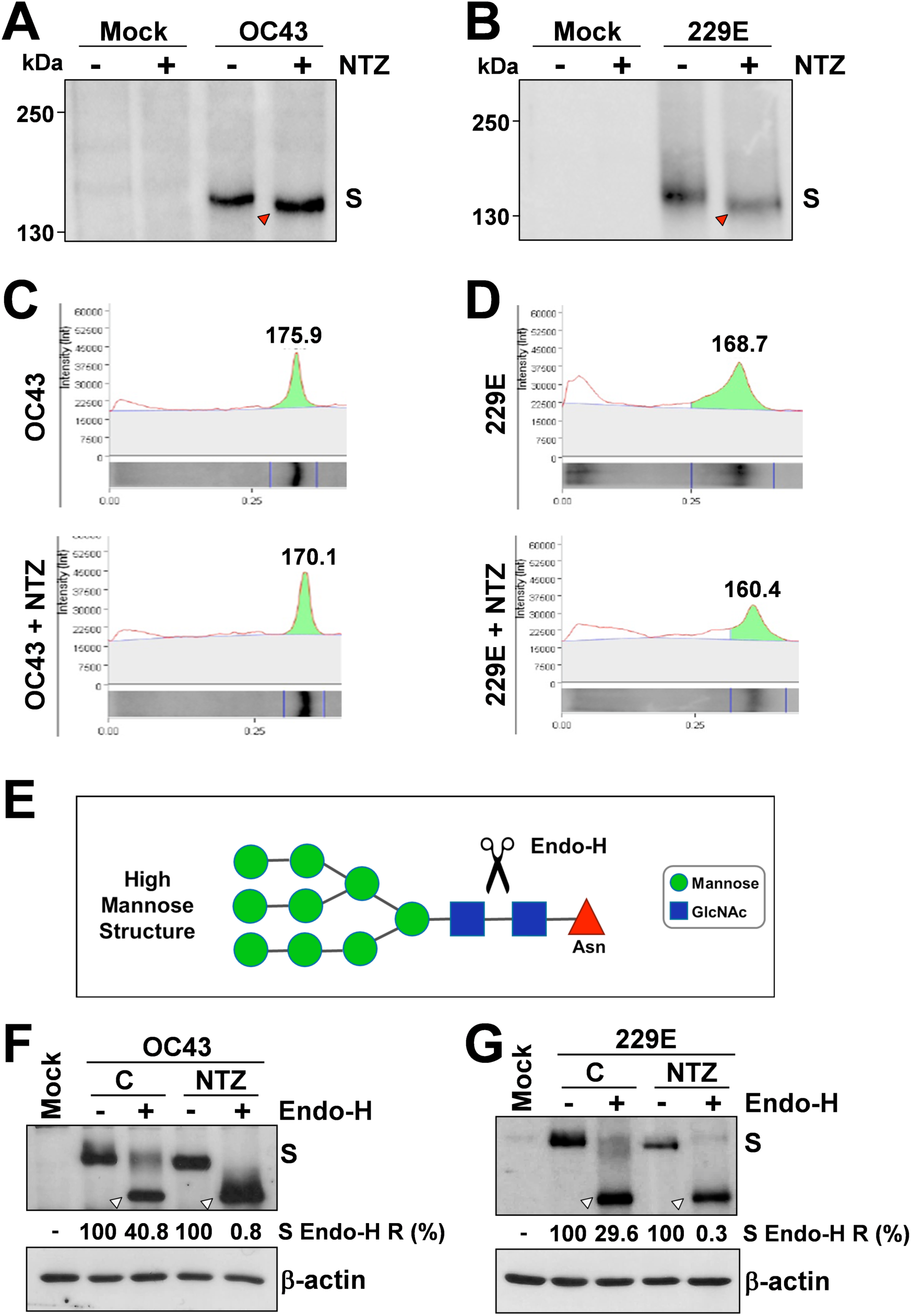
Nitazoxanide impairs HCoV-OC43 and HCoV-229E spike maturation at an Endo-H sensitive stage. (**A-D**) MRC-5 cells were mock-infected or infected with HCoV-OC43 (A,C) or HCoV-229E (B,D) at an MOI of 0.5 TCID_50_/cell and treated with NTZ (2.5 µg/ml) or vehicle immediately after the virus adsorption period. After 24h, WCE were analyzed for levels of S protein by IB using anti-S HCoV-OC43 (A,C) or anti-S HCoV-229E (B,D) antibodies. Red arrowheads indicate the faster-migrating forms of the HCoV-OC43 (A) and HCoV-229E (B) spike proteins in NTZ-treated cells. Spike proteins densitometric and molecular weight analysis (C,D) are shown. (**E**) Diagram of substrate specificity of endoglycosidase-H (Endo-H). Mannose (green circles), *N*-acetylglucosamine (GlcNAc, blue squares) and asparagine (Asn, red triangle) residues are shown. Scissors represent the cleavage site of Endo-H. (**F,G**) MRC-5 cells were mock-infected or infected with HCoV-OC43 (F) or HCoV-229E (G) at an MOI of 0.5 TCID_50_/cell and treated with NTZ (1µg/ml) or vehicle immediately after the virus adsorption period. After 24h proteins were digested with Endo-H (+) or left untreated (-) and processed for IB analysis using anti-S HCoV-OC43 (F) or anti-S HCoV-229E (G) antibodies, and quantitated by scanning densitometry. The Endo-H-cleaved faster-migrating S forms are indicated by white arrowheads. The percentage of Endo-H-resistant (Endo-H R) S protein in the different samples is indicated.

As indicated in the Introduction, glycosylation of coronavirus S proteins, like other cell surface glycoproteins, is initiated in the endoplasmic reticulum, adding the “high mannose” oligosaccharides; the mannose-rich sugar component is processed in the Golgi apparatus, and terminal glycosylation occurs in the *trans* cisternae of the Golgi apparatus (Xu and Ng, 2015).

To obtain insights on the effect of NTZ on HCoV S maturation, we therefore investigated whether nitazoxanide could affect HCoV-OC43 and HCoV-229E spike proteins terminal glycosylation. MRC-5 cells were infected with HCoV-OC43 or HCoV-229E (0.5 TCID_50_/cell) and treated with 1 μg/ml NTZ or vehicle starting immediately after the adsorption period. At 24h p.i., protein aliquots (20 μg) from NTZ-treated or control cells were subjected to digestion with Endo-H and then analyzed by immunoblot. The results, shown in Fig. 5F,G, indicate that, at this time, a fraction (approximately 40% in the case of HCoV-OC43, and 30% for HCoV-229E) of the spike proteins was found to be terminally glycosylated becoming Endo-H-resistant in control cells under the conditions described; notably, the spike proteins from NTZ-treated cells remained instead sensitive to digestion with the glycosidase up to 24h after synthesis. Because acquisition of Endo-H resistance is a marker for transport into the *cis* and *middle* Golgi compartments (Ohuchi et al., 1997), these results indicate that nitazoxanide may impair S protein trafficking between the ER and the Golgi complex.

## 4. Discussion

*Coronaviridae* is recognized as one of the most rapidly evolving virus family as a consequence of the high genomic nucleotide substitution rates and recombination (Cui et al., 2019). As indicated in the Introduction, to date, seven human CoVs have been identified, namely HCoV-229E, HCoV-NL63, HCoV-OC43 and HCoV-HKU1, globally circulating in the human population, and the highly pathogenic SARS-CoV, MERS-CoV and SARS-CoV-2. Among these, SARS-CoV-2 is responsible for a devastating pandemic that is causing unprecedented public health interventions (COVID-19 Excess Mortality Collaborator, 2022). Given the proportion of the COVID-19 pandemic, major efforts have been directed over the past months towards a global vaccination plan (https://cdn.who.int/media/docs/default-source/immunization/covid-19/strategy-to-achieve-global-covid-19-vaccination-by-mid-2022.pdf?sfvrsn=5a68433c_5). At the same time, the emergence of several SARS-CoV-2 spike variants that facilitate virus spread and may affect the efficacy of recently developed vaccines (Dong et al., 2021; Harvey et al., 2021; Planas et al., 2021; Carabelli et al., 2023), together with the short-lasting protective immunity typical of HCoV (Edridge et al., 2020), creates great concern and highlights the importance of identifying antiviral drugs to reduce coronavirus-related morbidity and mortality. So far, for SARS-CoV-2, two different RNA-dependent RNA polymerase (RdRp) inhibitors remdesivir (Santoro and Carafoli, 2020) and molnupiravir (Jayk Bernal et al., 2022; Wahl et al., 2021), and a viral protease inhibitor, Paxlovid (SARS-CoV-2 3CL protease inhibitor nirmatrelvir co-packaged with ritonavir), have been approved by health authorities in different countries (Hammond et al., 2022; Wen et al., 2022). On the other hand, no specific antiviral drug or vaccine are presently available for seasonal coronavirus infections.

Nitazoxanide has been proven to have a broad-spectrum antiviral activity (Li and De Clercq, 2020; Rossignol, 2014). In particular, nitazoxanide, its active metabolite tizoxanide and second generation thiazolides were found to be effective against several widespread RNA pathogens, including rotavirus, hepatitis C, and influenza and parainfluenza viruses in laboratory settings (Belardo et al., 2015; Korba et al., 2008; La Frazia et al., 2013, 2018; Rossignol et al., 2009; Piacentini et al., 2018; Stachulski et al., 2021), as well as in clinical studies (Haffizulla et al., 2014; Rossignol et al., 2006, 2009). In the case of *Coronaviridae*, we first reported the effect of the drug against a canine strain of the virus (CCoV S-378) in canine A72 cells in 2007 (Santoro et al., 2007). In 2015, Cao et al. showed that among 727 compounds tested against various strains of coronavirus, nitazoxanide was among the three most effective compounds tested (Cao et al., 2015). Nitazoxanide and tizoxanide were also found to be effective against MERS-CoV in LLC-MK2 cells with IC_50_s of 0.92 and 0.83 µg/ml respectively (Rossignol, 2016). As for SARS-CoV-2, at an early stage of the pandemic, Wang et al. reported that nitazoxanide inhibits SARS-CoV-2 replication in Vero E6 cells at low-μM concentrations (EC_50_  =  2.12  µM) (Wang et al., 2020); these observations were recently confirmed in different types of cells, including human lung-derived Calu-3 cells (Rocco et al., 2021; Son et al., 2022), as well as in animal models (Miorin et al., 2022). Tizoxanide was also recently found to be effective against SARS-CoV-2 in Vero E6 cells with an EC_50_ of 0.8 µg/ml (Riccio et al., 2022). More importantly several studies have recently shown an antiviral activity and clinical benefits of nitazoxanide in COVID-19 patients (Blum et al., 2021; Rocco et al., 2021; Rossignol et al., 2022; Silva et al., 2021). It should be pointed out that treatment of subjects with laboratory-confirmed seasonal HCoV infection with nitazoxanide administered orally twice daily for five days was also associated with improvement in time to return to usual health and time until subjects are able to perform normal activities (Rossignol et al., unpublished results). However, the mechanism at the basis of the antiviral activity of nitazoxanide against coronaviruses is not yet understood.

We now show that nitazoxanide has a potent antiviral activity against three human endemic coronaviruses, the Alpha-coronaviruses HCoV-229E and HCoV-NL63, and the Beta-coronavirus HCoV-OC43 in cell culture with IC_50_ ranging between 0.05 and 0.15 μg/ml and high (>330) selectivity indexes. The fourth known seasonal HCoV, HKU1, was not investigated because of its poor ability to grow in cell culture.

We found that nitazoxanide does not affect virus adsorption, entry or uncoating of HCoVs, but acts at postentry level, interfering with the spike S-glycoprotein maturation at concentrations that do not inhibit S and N protein expression in the infected cell. These results confirm in two different actively-replicating HCoV models our previous observation that nitazoxanide affects the maturation of the SARS-CoV-2 S glycoprotein in human cells transfected with plasmids encoding different SARS-CoV-2 spike variants (Riccio et al., 2022). The results are also in line with our previous studies on influenza and parainfluenza viruses, where nitazoxanide was shown to impair terminal glycosylation and intracellular trafficking of the class-I viral fusion glycoproteins influenza hemagglutinin and paramyxovirus fusion proteins (La Frazia et al., 2018; Piacentini et al., 2018; Rossignol et al., 2009).

As previously observed in the case of the hemagglutinin protein during human and avian influenza virus infection (La Frazia et al., 2018; Rossignol et al., 2009), and of exogenously expressed SARS-CoV-2 S protein in human cells (Riccio et al., 2022), nitazoxanide was found to hamper the spike protein maturation at an Endo-H-sensitive stage, thus preventing its final processing. This effect has been previously associated to the drug-mediated inhibition of ERp57, an ER-resident glycoprotein-specific thiol-oxidoreductase which is essential for correct disulfide-bond architecture of selected viral proteins (Piacentini et al., 2018).

Because of the critical role of the spike protein in coronavirus assembly (Fung and Liu, 2019; Santopolo et al., 2021a), hampering S maturation may result in hindering progeny virus particle formation; however, we cannot exclude the existence of additional mechanisms that may contribute to the antiviral activity of thiazolides (Hammad et al., 2022; Jasenosky et al., 2019; Rossignol, 2014).

As indicated in the Introduction, HCoV-229E, HCoV-OC43 and HCoV-NL63, as well as HCoV-HKU1, are distributed globally, and generally cause mild upper respiratory tract diseases in adults; however, they may sometimes cause life-threatening diseases in a subset of patients (Arbour et al., 2000; Chiu et al., 2005; Gorse et al., 2009; Jacomy et al., 2006; Lim et al., 2016; Liu et al., 2021; Morfopolou et al., 2016; Pene et al., 2003; Risku et al., 2010; Zhang et al., 2022). Interestingly, whereas HCoV-229E was suggested to be involved in the development of Kawasaki disease (Esper et al., 2005; Shirato et al.2014), HCoV-OC43 has been shown to have neuroinvasive properties and to cause encephalitis in animal models (reviewed in Bergmann et al., 2006; Cheng et al., 2020). Moreover, both HCoV-OC43 and HCoV-229E were shown to establish persistent infections in cell cultures (Arbour et al., 1999a,1999b), while the presence of HCoV-OC43 RNA was detected in human brain autopsy samples from multiple sclerosis patients (Arbour et al., 2000; Cheng et al., 2020).

These observations, together with the knowledge that HCoV infection does not induce long-lasting protective immunity (Edridge et al., 2020), highlight the need for broad-spectrum anti-coronavirus drugs. The results described in the present study indicate that nitazoxanide, which has been used for decades in medical practice as a safe and effective antiprotozoal drug (Rossignol et al., 2001, 2006, 2014), due to its broad-spectrum anti-coronavirus activity, may represent a readily available useful tool in the treatment of seasonal coronavirus infections.

## Supporting information

Piacentini et al. Supplementary Information

## Author Contributions

S. Piacentini, A. Riccio, SS and S. Pauciullo performed the study on the antiviral activity and the transfection experiments; S. Piacentini, A. Riccio, A. Rossi and SLF performed the analysis of protein synthesis and maturation. MGS and JFR designed the study; MGS wrote the manuscript. All authors contributed to the interpretation of the data and approve the content of the manuscript.

## Acknowledgments

The authors thank Lia van der Hoek (Academic Medical Center, University of Amsterdam) for providing the HCoV-NL63 Amsterdam-1 strain and the LLC-MK2 cell line. We also thank Dr. Elena Romano (University of Rome Tor Vergata) for assistance with confocal microscopy.

