## Supplementary material for "The FDA-approved drug nitazoxanide is a potent inhibitor of human seasonal coronaviruses acting at postentry level: effect on the viral spike glycoprotein": Piacentini et al. Supplementary Information

### Supplementary Figure Legends

**Supplementary Figure 1. Antiviral activity of nitazoxanide in human lung cells at different times after infection with OC43 and 229E HCoVs.**

(**A,B**) MRC-5 cells mock-infected or infected with HCoV-229E (A) or HCoV-OC43 (B) at an MOI of 0.1 $TCID_{50}$/cell were treated with 1 μg/ml NTZ (filled bars), or vehicle (empty bars) immediately after the adsorption period. Extracellular viral RNA levels were determined in the cell supernatants at 48h after infection by qRT-PCR. (**C,D**) MRC-5 cells mock-infected or infected with HCoV-229E (C) or HCoV-OC43 (D) at an MOI of 0.5 $TCID_{50}$/cell were treated as in (A,B). Extracellular viral RNA levels were determined at 24h p.i. by qRT-PCR. (**A-D**) Data, expressed as percent of untreated control, represent the mean ± S.D. of duplicate samples. *= $p < 0.01$; Student's *t*-test.

**Supplementary Figure 2. Effect of nitazoxanide treatment started at different times after infection with HCoV-229E in human lung cells.**

MRC-5 cells mock infected or infected with HCoV-229E at an MOI of 0.5 $TCID_{50}$/cell were treated with NTZ (1 μg/ml) or vehicle at the indicated times after virus adsorption. Virus yields were determined at 24h p.i. by infectivity assay. Data, expressed as $TCID_{50}$/ml, represent the mean ± S.D. of duplicate samples. *= $p < 0.01$; Student's *t*-test.

**Supplementary Figure 3. Nitazoxanide does not inhibit HCoV-229E spike protein expression in MRC-5 lung cells.**

Confocal images of HCoV-229E spike glycoprotein (red) in MRC-5 cells mock-infected or infected with HCoV-229E at an MOI of 0.5 $TCID_{50}$/cell and treated with NTZ (1 μg/ml) or vehicle for 24h. Nuclei are stained with Hoechst (blue). Merge images are shown. Scale bar, 20 μm.

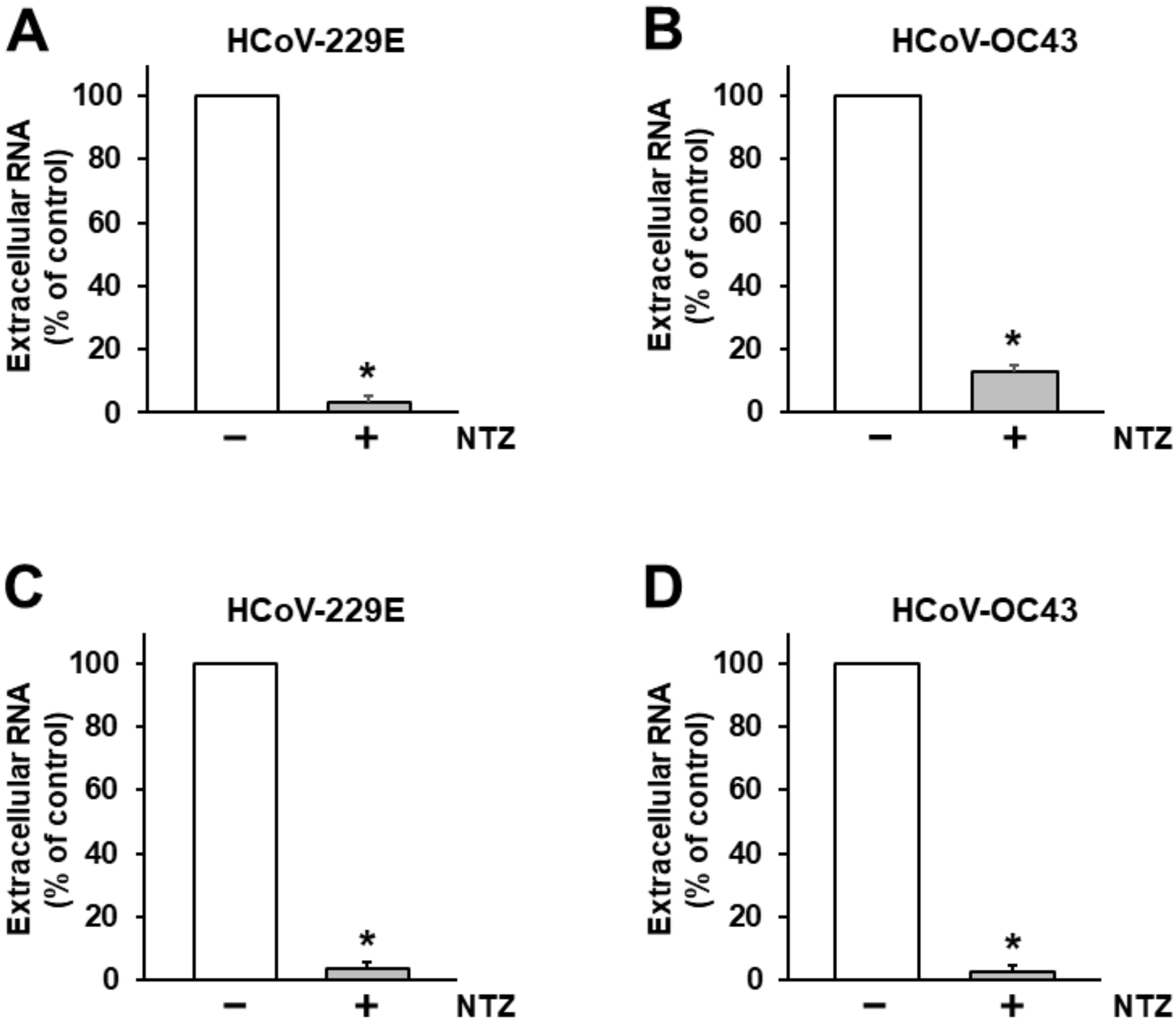
A
HCoV-229E
Extracellular RNA (% of control)
100
80
60
40
20
0
*
−
+
NTZ
B
HCoV-OC43
Extracellular RNA (% of control)
100
80
60
40
20
0
*
−
+
NTZ
C
HCoV-229E
Extracellular RNA (% of control)
100
80
60
40
20
0
*
−
+
NTZ
D
HCoV-OC43
Extracellular RNA (% of control)
100
80
60
40
20
0
*
−
+
NTZ

**Figure S1**

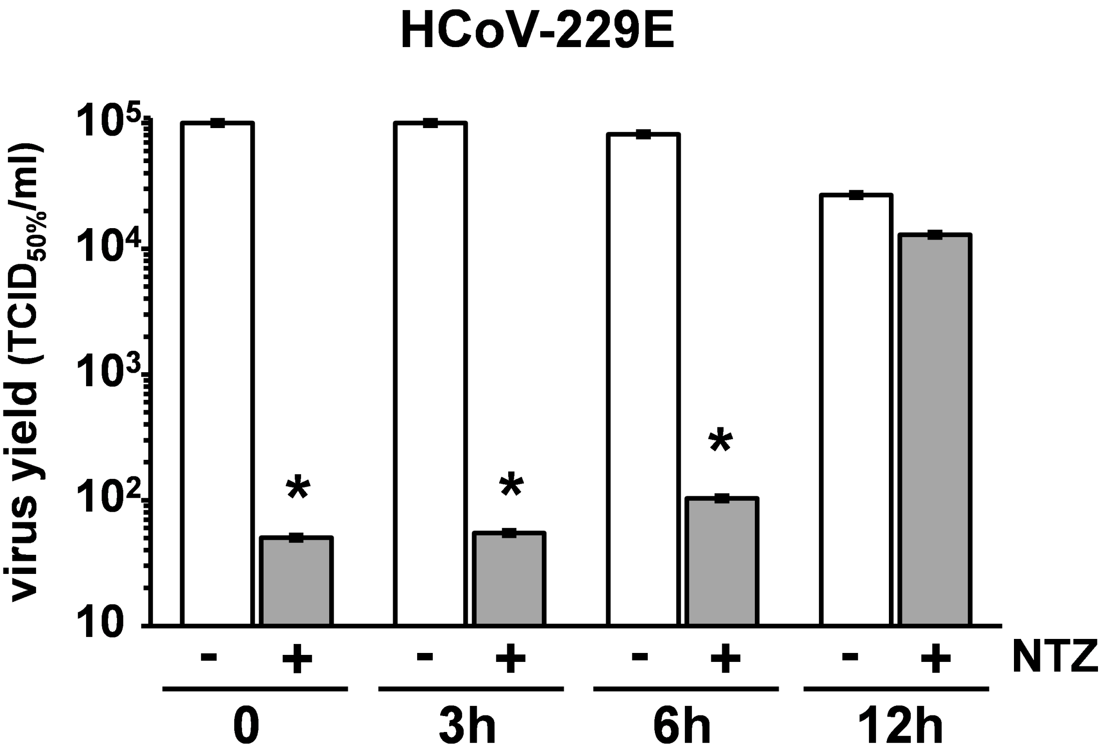
HCoV-229E
virus yield (TCID50%/ml)
10
10²
10³
10⁴
10⁵
*
*
*
- + - + - + - + NTZ
0
3h
6h
12h

**Figure S2**

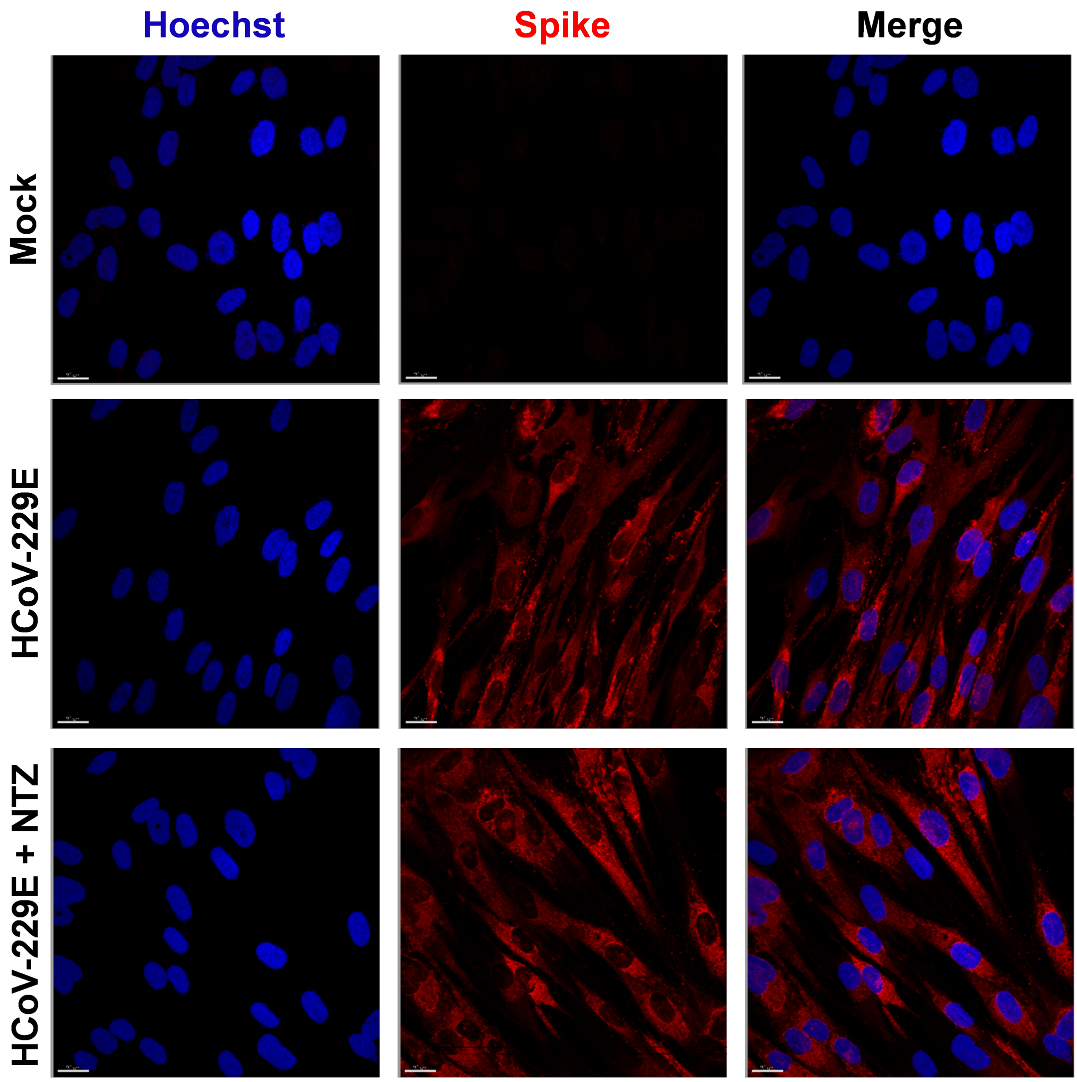


Figure S3
